# Electrophysiological correlates of lucid dreaming: sensor and source level signatures

**DOI:** 10.1101/2024.04.09.588765

**Authors:** Çağatay Demirel, Jarrod Gott, Kristoffer Appel, Katharina Lüth, Christian Fischer, Cecilia Raffaelli, Britta Westner, Xinlin Wang, Zsófia Zavecz, Axel Steiger, Daniel Erlacher, Stephen LaBerge, Sérgio Mota-Rolim, Sidarta Ribeiro, Marcel Zeising, Nico Adelhöfer, Martin Dresler

## Abstract

Lucid dreaming (LD) is a state of conscious awareness of the current dream state, predominantly associated with REM sleep. Research progress in uncovering the neurobiological basis of LD has been hindered by low sample sizes, diverse EEG setups, and specific artifact issues like saccadic eye movements and signal non-stationarity. To address these matters, we developed a multi-stage preprocessing pipeline that integrates standardized early-stage preprocessing, artifact subspace reconstruction, and signal-space projection. This approach enhanced data quality by precisely removing saccadic potential effects even in setups with minimal channels. To robustly identify the electrophysiological correlates of LD, we applied this methodology to LD data collected across laboratories (pooled N = 44) and explored sensor-and source-level markers hypothesized to underlie LD. Compared to non-lucid REM sleep, we observed few robust differences on the EEG sensor level, which is in line with recent findings. In contrast, on the source level, gamma1 power (30-36 Hz) showed increases during LD in left-hemispheric temporal areas, which might reflect verbal insight processes. Gamma1 power also increased around the onset of LD eye signaling in right temporo-occipital regions including the right precuneus, in line with its involvement in self-referential thinking. Reductions in beta power (12-30 Hz) during LD in right central and parietal areas including the temporo-parietal junction are potentially associated with a conscious reassessment of the veridicality of the currently perceived reality. Notably, functional connectivity in alpha band (8-12 Hz) increased during LD, in contrast to the reductions typically seen in psychedelic states, highlighting enhanced self-awareness. Taken together, these findings illuminate the electrophysiological correlates of LD state, and may serve as a basis to uncover neural mechanisms at the time point of lucid dream insight.

## Introduction

During lucid dreaming (LD) subjects know they are dreaming, and sometimes are able to control the oneiric content. Lucid dreamers are aware of the hallucinatory character of ongoing perceptions while asleep (1–5), which often allows them to voluntarily interact with their internal world models (6) via fully immersive simulations (2, 5). LD happens predominantly during rapid-eye-movement (REM) sleep, but can also occur during sleep onset (N1) and light sleep (N2) stages (7–10).

Electroencephalographic (EEG) research of LD was initiated in the late 1970s with the introduction of the eye signaling technique: experienced lucid dreamers were instructed to perform intentional left-right-left-right movements in their dreams (11–13), which in turn can be objectively assessed via electrooculography (EOG), as eye muscles are not affected by the general muscle atonia experienced during REM sleep (14, 15). This technique has since become the gold standard method in LD research, allowing for electrophysiological recordings of LD in multiple studies by different research groups (16).

Despite five decades of EEG research on the topic, these studies still paint an inconclusive picture of LD neurophysiology (17, 18): earlier findings such as increased occipital alpha (8-12 Hz) (19) parietal beta (13-19 Hz) (20) or frontal gamma (36-45 Hz) underlying LD (21) could not be substantiated after adequate removal of non-neural artifacts (18). One independently replicated finding is that reduction in delta-band (0.5-2.9 Hz (22); 2-4 Hz (18)) activity differentiates LD from non-lucid dreaming, potentially pointing to alterations in sleep depth.

LD is a learnable skill (23, 24), however still cannot be reliably induced (25), leading to very small sample sizes in most studies. The rarity of LD (26, 27) and thus lack of statistical power in the typical non-representative studies may be overcome by aggregating data from multiple studies or recording sites, which however requires considerable effort in harmonization of datasets. Here we developed an adaptive preprocessing pipeline for valid EEG data aggregation tailored to LD, addressing the field’s inherent confounds such as saccadic artifacts and assumptions of signal non-stationarity in distinct EEG layouts, which we detail next.

A particular concern in LD recordings are saccadic artifacts: higher eye movement density during LD (28) will be reflected in frontal potentials if not properly accounted for (17). A recent study demonstrated (18) that a unique signature of LD found in an earlier study (21), namely fronto-lateral gamma at 40 Hz activity, could be explained by improper cleaning of saccadic potentials (SPs), which are resistant against traditional, regression-based cleaning approaches (18, 29). SPs are minor spike deflections in the horizontal EOG (18, 29, 30) and noticeable just before saccadic eye movements, especially in the direction opposite to the saccade’s trajectory. While independent component analysis (ICA) has been shown to clean SPs that happen during LD (18), ICA may overclean data when applied to few EEG channels (31). As an alternative, we provide a preprocessing solution that works for low-density EEG setups and hence is applicable for analyzing data from wearable devices intended for home use (32) as well as polysomnographic recordings in the laboratory.

Our proposed preprocessing pipeline further deals with signal nonstationarities that are likely to be encountered within the cognitively complex computations of the LD state (33). Note that not just sleep per se (14), but REM sleep in particular is inherently non-stationary, as it can be subdivided into phasic and tonic periods (34) that have distinct spectral signatures (35). Many established preprocessing tools such as ICA often assume stationary signals (36), and hence might overclean data when spectral aspects are dynamically shifting in a non-stationary manner. Again, our specifically designed preprocessing protocol is set up to deal with this issue.

After the uniformization of preprocessing steps across different EEG setups, we aimed to investigate the spectral correlates and functional connectivity patterns across all traditional frequency bands, as well as measures of entropy, complexity and fractal dimension to discern how LD differs from both REM sleep and waking state. Further, we also explore the dynamic nature of evolving LD over time. Specifically, it was argued that the moment of lucid insight might be more characteristic of LD than any tonic marker (17). We therefore explore the neural dynamics that occur in the vicinity of the consciously performed eye movements that first signal LD. Figure 1B depicts an overview of the preprocessing and analysis protocol.

**Figure 1.**
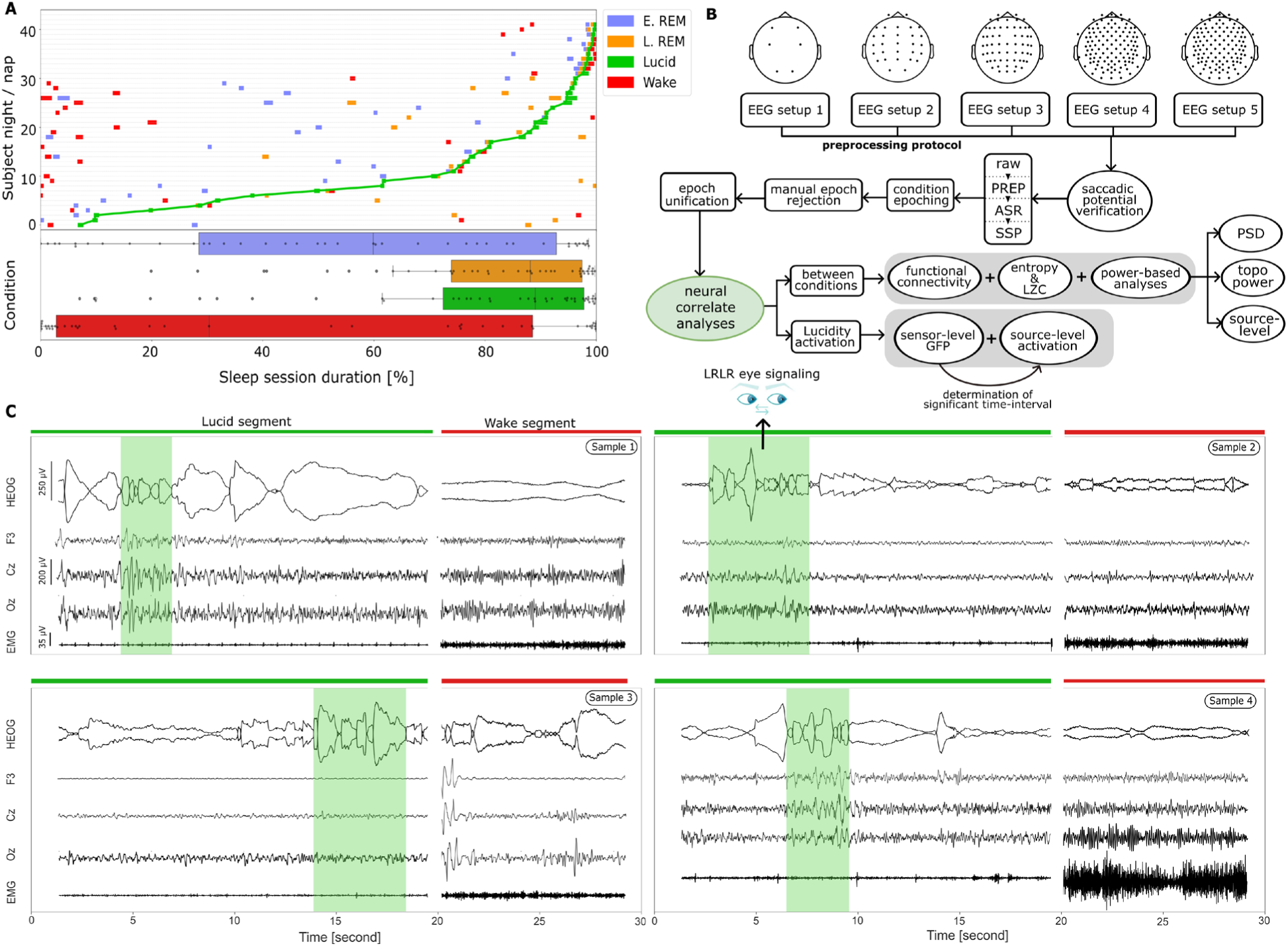
**A:** Time table of conditions over sessions. The upper section shows the individual distributions of condition on-and offsets (colored bars) with respect to a standardized timeline, where 0% denotes recording onset and 100% recording offset. Individual recordings are sorted along the y axis with regards to LD onset, showing how most LD episodes are collected towards the end of a recording session. Also note that waking condition intervals occur early in the timeline as this was the main selection criterion for this condition (see text for details). The lower section depicts the distribution of each condition segment’s center over the standardized timeline using box plots and individual data points. A notable gap between early and late REM states, accounting for about 50% of total sleep duration, indicates the passage of one or more sleep cycles during both nap and full-night experiments. The LD segment is positioned close to the later REM phase, with selection slightly earlier on average. **B:** Overview of the EEG pre-processing, post-processing and analysis protocol employed in this study. Note that this protocol can handle and integrate diverse EEG setups to allow robust neural correlation analyses. **C:** Illustration of amplitude and waveform characteristics of EEG, EOG, and EMG activities between lucid dreaming and wakeful states, showcasing representative segments from four participants. LD: lucid dreaming; REM: rapid eye movement sleep; PREP: early-stage EEG processing pipeline; ASR: artifact subspace reconstruction; SSP: signal space projection; PSD: power source density; topo: topography.

Importantly, our study pioneers the use of electrophysiological source localization in LD, leveraging high-density sensor data to uncover the neural correlates of the LD state through spectral analysis of neuroanatomical structures. Given the inconclusive state of the field so far, we approached our analyses without specific hypotheses about spectral bands or brain regions affected, thus providing a comprehensive exploration of the electrophysiological correlates of LD.

## Methods

### Participants

The intrinsic rarity of LD calls for a collaboration across laboratories to gather a sample of adequate size. Previously unpublished LD data was gathered from multiple laboratories: the Donders Center for Cognitive Neuroimaging at Radboud University, Netherlands (RU, 12 recordings), University of Osnabrück, Germany (UO, 5 recordings), Max Planck Institute, Munich, Germany (MPI, 10 recordings), Brain Institute at Federal University of Rio Grande do Norte, Brazil (UFRN, 2 recordings) and Department of Psychology, Stanford University, United States (27 recordings). We obtained signed informed consent prior to recording from all subjects, and ethical approval was acquired from the corresponding university institutional review boards (approval codes CMO 2014/288, 4/71043.5, 094-13, CEP-UFRN, and 061/2008) for RU, UO, MPI, and UFRN respectively). Details regarding the dataset from SU can be reached in the previous study (37). We excluded data with less than 6 EEG channels, continuous eye signaling during LD, unclear lucidity eye signals, having no possibility to capture two distinct REM conditions and containing persistent noise that we could not clean up. This led to the exclusion of 16 recordings, with the final sample consisting of 26 subjects (25.0 ± 5.1 years, 20 females) familiar with LD experiences. Of those, five subjects contributed two nap and night sessions each, one contributed three nap sessions and the other one subject contributed thirteen nights, yielding a sample size of 44 sleep recordings: 5 recordings consist of 6 channels, 20 recordings consist of 29 channels, and the remaining 19 are high-density recordings with 64 or 128 channels.

### Data recording

Polysomnography recordings started when participants went to bed. Bed times differed across labs: With regards to data collected at RU, participants went to bed in the morning hours after slight sleep deprivation, whereas at UO, LMU, UFRN and SU, full nights were recorded. For this, we used different EEG devices and setups, namely: at UO, a 6-channel Somnoscreen PSG device utilizing a 10/20 layout (F3, F4, C3, C4, O1, O2); at RU, a 64-channel actiCAP active electrode EEG system (Brain Products) with a 10/10 layout; at MPI, a 128-channel easyCAP passive electrode EEG system (Brain Products) featuring a 10/05 layout; at UFRN, a 32-channel EMSA BNT-36 passive electrode EEG system with a 10/20 layout; and at SU, a 32-channel passive electrode system (Neuroscan, El Paso, TX, USA) containing 29 EEG electrodes placed in a 10/20 layout. After each sleep session, free reports of the last remembered dream were collected via audio recordings and later transcribed by researchers. The recordings have bit depths of either 8-bit or 16-bit, varying across the five different datasets.

### Task

Before all sleep sessions, we instructed participants to move their eyes to the left, then to the right, and repeat this sequence at least once as soon as they became aware that they were currently dreaming. If successful, this well-established maneuver yielded a so-called LRLR signal during sleep (16) which can be easily discerned in horizontal EOG recordings (see Figure 1C). Beyond this common task, protocols varied between subjects and labs, encompassing flying, counting, two-way communication, motor movement (hand-clenching), and mind-wandering tasks. Given the diversity of event protocols across datasets during lucid dreaming, there is a wide range of eye signaling patterns, from initial and final LRLR sequences to multiple intermediate eye movements that mark the beginning and end of specific events. The duration of these eye signals also varies, ranging from brief movements signaling event onsets to longer eye movements that are considered voluntary ocular events. Figure 2 illustrates EEG segments that capture LD episodes contrasted with wakefulness across all recordings.

**Figure 2.**
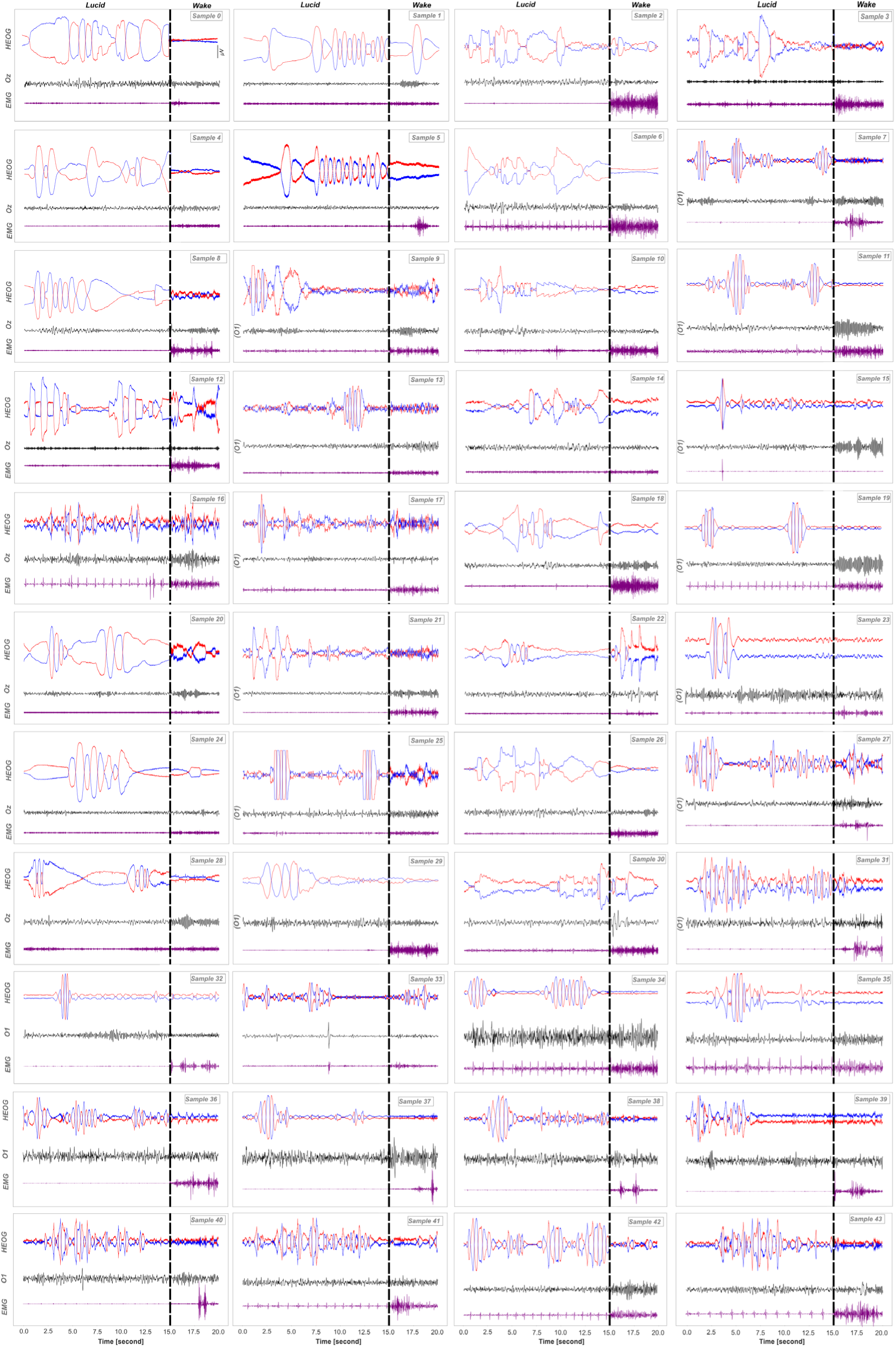
Single-subject horizontal EOG, EEG and EMG timeseries during eye signaling within lucid dreaming and waking. Either the Oz or O1 EEG channel is shown, with O1 selected when Oz is not available.

### Sleep scoring

All sleep data were scored either by at least one human expert or by the U-Sleep v2.0 (38) automated model, using 24 sparse EEG channels along with an additional EOG channel. We trained human raters to score the data according to the criteria of the second version of the sleep scoring manual by the American Association of Sleep Medicine (AASM). Accordingly, we labeled continuous polysomnographic recordings within each 30-second non-overlapping interval. Sleep scoring was done blind with no auxiliary information other than the PSG data. In cases when sleep score data of only one human rater was available, it was supplemented with ratings by an automatically trained model, namely YASA (39). This yielded two scores for each recorded session (note that epochs with a YASA confidence rating of below 80% were rejected from analysis).

### Selection of conditions

REM and waking segments were chosen for quantitative comparisons with LD segments within the same sleep session. We selected time intervals from each session recording that matched the duration of lucid REM segments for comparison with the states of waking, non-lucid early REM (E. REM), and non-lucid later REM (L. REM). Two such temporally distant REM intervals were selected to test the stability of potential LD correlates against confounding effects of night time (e.g. due to changes in sleep pressure (40)). First the data was divided into 30-second intervals, and aligned with sleep score intervals. For each channel and time interval, we averaged the computed standard deviations of time samples to derive a singular value per interval, excluding those below the 50th percentile to lessen the impact of motion artifacts. We selected waking and REM sleep intervals matching LD segment lengths, focusing on ensuring E. REM and L. REM states were ideally a sleep cycle apart or separated by other sleep stages by aiming low temporal proximity. This simplified approach aimed to maintain the natural spacing between early and later REM phases for accurate comparison, prioritizing intervals that offered clear distinctions in REM sleep stability and duration.

We concluded a visual inspection of the selected segments in the temporal domain to ensure no significant artifacts were included. A schematic overview of the project including the visual depiction of conditions for each subject with the overall distributions across normalized sleep duration is shown in Figure 1. Note that for the waking condition, we primarily selected data at the beginning of the experiment, specifically during periods when the subjects’ eyes were closed. This strategy aimed at choosing a waking state where subjects were relaxed, not conversing, and free from physically induced artifacts. Ensuring the waking represented a relaxed state was crucial for the accuracy of our analysis.

### Multi-stage preprocessing protocol

Our newly designed multi-stage preprocessing protocol specifically mitigates saccadic artifacts and signal non-stationarity, which are potential confounds that are especially important to consider in electrophysiological measures of LD. Saccadic potential (SP) artifacts are particularly prevalent and problematic in the context of LD research due to the eye movements characteristic of this state. The ocular movements during LD are both involuntary (18) -the rapid eye movements that define REM sleep, and voluntary by instructing the subjects to perform eye signals in order to be able to determine their lucidity state and subsequent events (16). A recent study (18) suggested that ICA successfully cleansed the SP artifacts during LD, but this was achieved with relatively high-density EEG data (32 channels). Importantly, ICA is sensitive to spatial sampling (31) and therefore does not provide robust cleaning for low-density variations (e.g. 6 channels). In this study, we have developed an alternative multi-stage preprocessing protocol that addresses non-stationarity of LD EEG and allows for inclusive analyses with low-density EEG setups (see see Figure 1B). The pipeline is adaptive given that with minimal parameter tuning effort, it offers semi-automatic signal cleaning in a data-driven manner. By doing so, we could standardize the data across diverse EEG setups (in both low and high density layout variations) and, in later stages, unify them to enable large-sample-size analyses of the neural correlates of LD.

After standardizing the channel labels across datasets and assigning channel types (EEG, EOG, EMG, ECG), we applied the three-stage preprocessing protocol consisting of the PREP pipeline (41), artifact subspace reconstruction (42) (ASR) and signal-space projection (43) (SSP), each taking unsegmented, continuous data as input in order to avoid condition-based biases. Note, however, that before running the algorithms, we cropped the data between the onset of the earliest condition (which most often was E. REM; see Figure 1A) and the offset of the latest condition (most often LD) in order to save computation time and focus on the signals of interest.

### Stage 1: PREP pipeline

The PREP pipeline offers standardized early-stage (pre-artifact rejection) EEG preprocessing ensuring consistency across diverse datasets (41). Apart from its robust line-noise removal capabilities, PREP effectively resolves a major circular problem of EEG re-referencing (identifying bad channels depends on a good average reference, while a good average reference pre-supposes bad channel exclusion), which is accomplished by iteratively detecting and excluding bad channels and thus refining the average reference with each iteration. We further improved bad channel detection in high-density setups via enabling the included RANSAC (random sample consensus) options which requires at least 19 channels. Prior to employing PREP, we applied a 1-49 Hz IIR bandpass filter to the data. For applying these functions in this study, we employed PyPREP (44) within a Python 3.9 environment.

### Stage 2: Artifact subspace reconstruction

Artifact Subspace Reconstruction (ASR), a non-destructive, dynamic method for cleaning multi-channel EEG artifacts, was selected for its data-driven, adaptive nature (42), that effectively managed the discrepancies from diverse artifact sources, and referencing setups in our dataset. Before applying this algorithm, we high-pass-filtered data at 2 Hz in accordance with (18) and following empirical recommendations for non-destructive EEG cleaning procedures (ASR (45) and independent component analysis (46–48)). ASR trains on a clean EEG segment to establish a baseline model of clean signals, which is then used to identify deviations in test data. Training data was selected as follows: First, we divided the data into 300-second consecutive intervals with 75% overlap. For each interval, standard deviations were calculated within each channel and summed across channels. The four intervals with lowest sums were then concatenated, yielding 20 minutes of training data for each sleep EEG recording. We chose to include multiple intervals in order to offer varying levels of stationarity for the training procedure, which should yield more robust and versatile results. Using sample covariance as an estimator, detected artifacts were cleaned within two-second intervals with 33% overlap. These parameters offer enough precision to clean artifacts in a sleep stage-specific manner. We decided against using a newer, Riemannian version of ASR (rASR (49)) for reasons of efficiency and visual inspection which revealed remarkably better cleaning of SPs in the standard version. ASR was applied using respective functions in the Python package meegkit within a Python 3.9 environment.

### Stage 3: Signal-space projection

While PREP and ASR handled the majority of artifacts resulting from SP and other miscellaneous ocular and heart rate-related activities (e.g. potentially increased heart rates due to motor performance during LD (50)), visual inspection of the data suggested they did not entirely address alls aspects of EOG and ECG artifacts (see Figure 3C). To resolve this problem we then used signal-space projection (SSP), a spatial filtering technique that leverages all available bio-potential sensors including EOG and ECG for non-destructive artifact cleaning. More specifically, it minimizes artifact interference in EEG data by utilizing EOG and ECG channels to identify distinct EOG and ECG artifact patterns, computing orthogonal projection vectors for these, and transforming the EEG data into a subspace where artifact influence is reduced. SSP was applied via respective functions in the MNE-Python toolbox (51).

**Figure 3.**
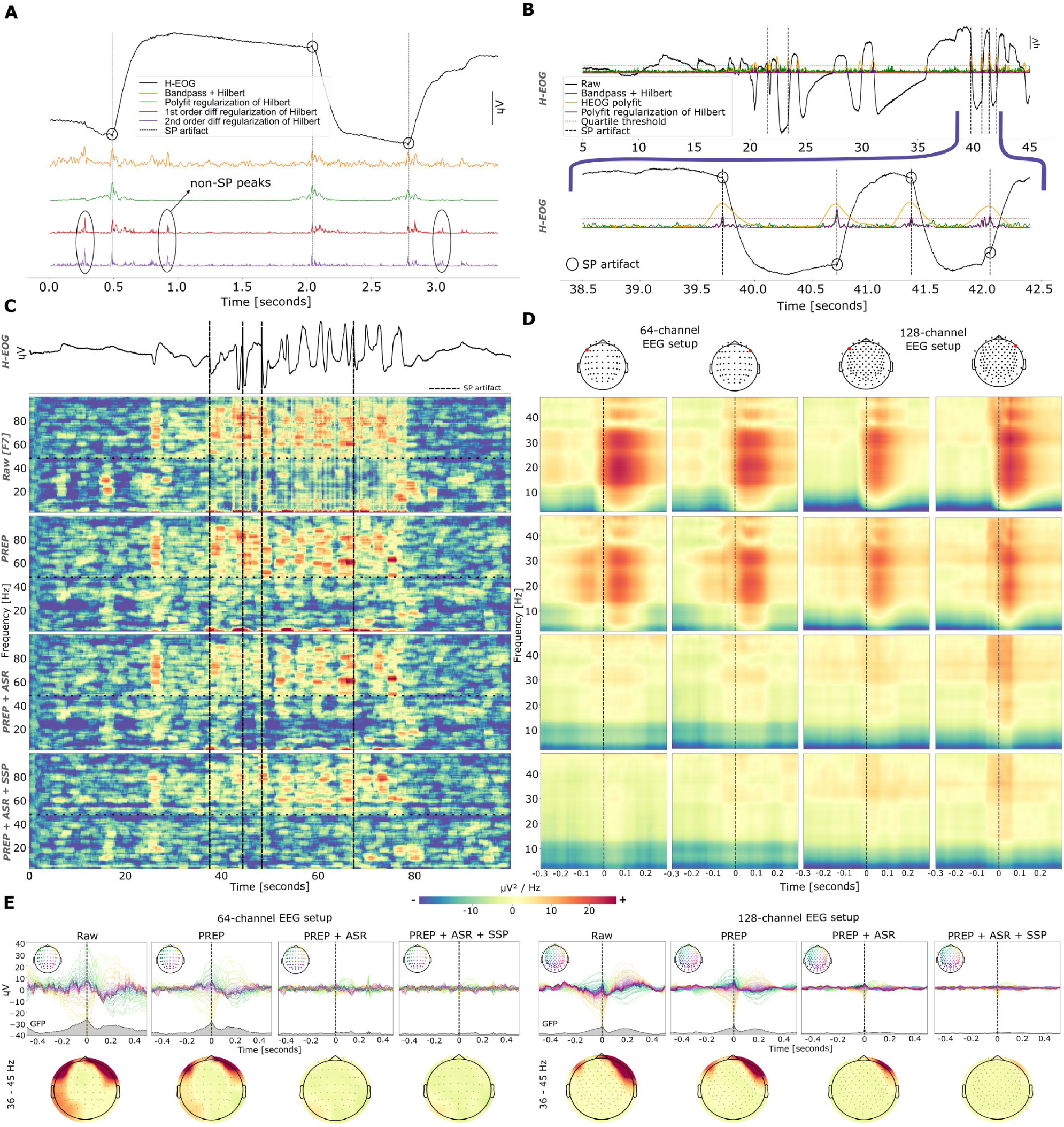
**A:** Example comparison of differently regularized Hilbert series of HEOG signals with regards to SP detection sensitivity and specificity. Note that derivative-based transforms are sensitive to non-SP-related activity as shown by the highlighted peaks. **B:** Thresholding of polynomial regularized Hilbert series of HEOG detects SP events. In this example, a 99.9 percentile was selected. **C:** Exemplary effects of SP events during LD on ERSP effects at channel F7. Better cleaning of broadband SP artifacts, especially below 48 Hz, is shown with each cumulative preprocessing step. **D:** Grand average ERSPs of SP events to demonstrate SP cleaning across cumulative preprocessing steps. The left two columns depict recordings from 64-channel active electrode setups (N = 16 SPs), the right two columns from 128-channel passive electrode setups (N = 63 SPs). The left and right column of each column pair show data from channels F7 and F8, respectively. **E**: Upper row: Grand average butterfly plots and GFPs around SPs for each cumulative preprocessing step. Bottom row: Topographies of SP artifacts for each cumulative preprocessing step in the ∼40 Hz band (36-45 Hz (27, 30)). Left panel: 64 channel active electrode setup. Right panel: 128 channel active electrode setup. A remarkable flattening of SP-related GFPs can be shown in all of these setups. HEOG: horizontal electro-oculogram. SP: saccadic potential. LD: lucid dreaming. ERSP: event-related spectral perturbation. GFP: global field power.

### Saccadic spike potential cleaning

Before evaluating the efficacy of our preprocessing steps with regards to cleaning SP artifacts, we first required a valid identification of SPs. For this, we investigated continuous 100-second HEOG data intervals from every night recording that included both initial and final LRLR signals, ensuring equal padding before the first and after the last signal. This selection was chosen to provide prominent eye movements that included LRLRs for later validation of SP detection. Several filters were applied to these HEOG data in order to test which of them best help identify time points of SPs. Specifically, we applied a band-pass filter between 30 Hz and 100 Hz as suggested previously (29) with a subsequent Hilbert transform; a Hilbert transform regularized with a 2nd-order polynomial fit; a 1st-order derivative; and a 2nd-order derivative. We chose the 1st-and 2nd-order derivatives and the 2nd-order polynomial fitting as regularization factors after Hilbert transform, normalizing the factor domains by dividing them by their maximum value, which scaled from 0 to 1. The polyfit regularization is less likely to get affected by other miscellaneous artifact-based peaks. This is because polyfit performs a slope approximation to regularize the Hilbert series by quantifying the negative or positive dumping factors that occur after the SP event.

Accordingly we selected a polyfit regularizer to find SP events. This was done via visual inspection, opting for conservative selection in order to retain potential signals of interest. Once this filter was selected, we manually selected a threshold percentage that most precisely identified SPs. Systematic differences in HEOG electrode placement between recordings from different datasets necessitated setting individual threshold values for each dataset. The maximum value of data above the threshold then pointed to the time point of putative SPs. Before further validation of our preprocessing/cleaning pipeline, we manually discarded false-positives (i.e. above-threshold filter peaks that are not associated with a saccade).

To evaluate the steps of our preprocessing pipeline with regards to cleaning SP artifacts, we investigated the F7 and F8 channels as a scalp position common to all datasets with high fronto-lateral ocular reflections. Event-related potentials were calculated for each cumulative step in the pipeline (i.e., PREP; PREP followed by ASR; PREP followed by ASR followed by SSP). In the time domain, we investigated Hilbert envelopes across all EEG channels as well as SP-related GFP fluctuations. In the time-frequency domain, we calculated event-related spectral perturbation (ERSP) with multitapers (frequency resolution: 0.1 Hz; time bandwidth value: 8). No logarithmic transformation was applied in order to preserve high contrast values. While our statistical analyses did not exceed 50 Hz, for this validation step we calculated frequencies up to 100 Hz for a more sensitive picture of the effects of our preprocessing steps.

### Epoching and post-processing

We segmented the cleaned subject data into 4-second windows with a 2-second overlap (i.e., 50% padding) for four conditions (E. REM, L. REM, LD, and waking). Despite our multi-stage preprocessing protocol largely eliminating SP artifacts, we conducted manual inspections of the epoched data for residual miscellaneous and movement-related artifacts, discarding contaminated epochs. We restricted all analyses regarding neural correlates of LD to the spectral domain between 2 and 48 Hz in order to avoid spectral leakage from bandpass filtering.

### Non-topographical global analyses

For the non-topographical analysis, we analyzed 6 common channels (F3, F4, C3, C4, O1, O2) from all 44 subject data, ensuring maximum sample size and consistent scalp position contributions. Since all other (topographical) analyses require higher spatial sampling, we selected a subset of 19 participants that share 59 channel positions, and limited our analysis to these 59 channels (see Table S1) in order to balance topographical detail with sample size and comparability. This decision necessitated the exclusion of electrodes ’TP10’, ’Iz’, ’FT10’, ’FT9’, and ’TP9’ from 64 channels EEG layout, as some subjects’ data did not include readings from these electrodes.

#### Power analysis

We selected epochs unified across different EEG setups. For each of the 6 channels and epochs, we extracted multitaper power spectral density (mPSD) values ranging from 2 to 48 Hz and applied a dB conversion (10 times the base-10 logarithm of the mPSD). We averaged the values across channels and epochs and extracted the 95% confidence bands for the conditions. Furthermore, we averaged PSD values within six different frequency ranges (delta: 2 -4 Hz; theta: 4-8 Hz; alpha: 8 - 12 Hz; beta: 12 - 30 Hz; gamma1: 30 - 36 Hz; gamma2: 36 - 45 Hz).

#### Entropy and complexity analysis

In accordance with previous research (52), we attempted to extract consciousness levels in EEG time-series via Lempel-Ziv complexity (LZc) (53) and two selected entropy markers—permutation entropy (PE) (54) and sample entropy (SE) (55). LZc is a non-linear dynamic measure that quantifies the rate of new pattern generation in a time series, making it a valuable tool for assessing the complexity of EEG signals, which are known for their non-linear and non-stationary nature (56). PE offers a robust and computationally efficient way to capture the order relations between values in a time series, providing insights into the dynamical changes in EEG data (54). We derived condition-specific estimates via averaging over the channel-specific values. The formulas of PE and SE are shown in Eq. 1 and Eq. 2, respectively.

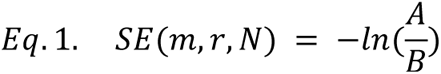

where m is the embedding dimension represents the length of the sequences being compared, *r* is the tolerance for accepting matches, *N* is the number of data points in the time-series, *A* is the number of matched pairs of sequences of length *m* + 1 and *B* is the number of matched pairs of sequences of length *m*.

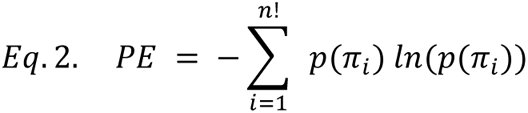

where *p*(*π*_*i*_) is the probability of a given permutation *π*_*ii*_ occurring within the time-series, and *n*! is the factorial of the embedding dimension *n*, representing the total number of possible permutations.

#### Fractal analysis

The Higuchi Fractal Dimension (HFD) (57) is a mathematical method used to quantify the fractal properties of time series data. HFD measures the complexity and self-similarity of a signal, providing a single value that represents the fractal dimension of the data. The HFD *D* of a time series *X* = {*x*(1), *x*(2), *x*(*n*)} is estimated in Eq. 3.

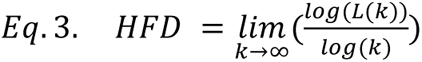

where *L*(*k*) is the average length of the time series for each scale *k*, given by:

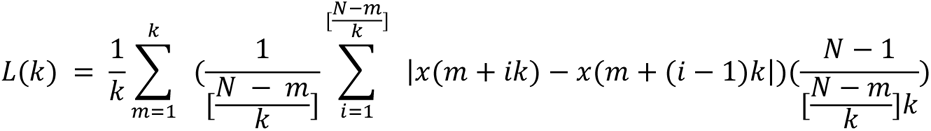

where the fractal dimension *D* is calculated by analyzing the complexity of the time series *X* over various scales *k*.

#### Sensor-level topographical power analysis

For each subject and condition, we computed multitaper power spectral densities (mPSD) across various frequency ranges. We derived subtracted power values from these contrasts and normalized each contrasted dataset across all channels, standardizing amplitudes by maximum division.

### Surface-based source reconstruction

We employed two variations of the Minimum Norm Estimate (MNE) method for surface-based source reconstruction: dynamic statistical parametric mapping (dSPM) (58) and exact Low-Resolution Electromagnetic Tomography (eLORETA) (59). We opted to employ both methods in a complementary manner. We recognized that dSPM, with its noise-normalization feature, excels in highlighting brain regions with significant activity by effectively reducing background noise. This makes it particularly useful for detecting subtle changes in brain activity across different conditions. On the other hand, eLORETA offers enhanced spatial accuracy in minimizing localization errors assuming a smooth distribution. Furthermore, applying both dSPM and eLORETA allowed us to cross-validate our findings, increasing our confidence in the robustness of our results.

The initial step in our pipeline involved applying average re-referencing to all subject datasets. For cortical mapping, we selected the ’fsaverage-ico-5-src.fif’ from the boundary element model (BEM) as our source space template, ensuring a high-resolution cortex representation. The forward solution was calculated with a minimum distance (mindist) parameter set to 5.0 mm minimum distance constraint. To calculate noise covariance, we developed an algorithm to identify baseline segments from the entire sleep dataset for each subject. This algorithm evaluated the standard deviation (STD) of 10-second windows, with an 80% overlap, across the raw data. The segment with the lowest STD, sufficiently distanced from all experimental conditions to avoid overlapping neural processes, was selected as the baseline. This baseline segment was then used to compute the noise covariance, which informed the creation of the inverse operator, applying a depth parameter of 0.8 to fine-tune source localization. For source power spectral density (PSD) estimation, we set the signal-to-noise ratio (SNR) parameter at 3, determining the lambda value for regularization. We utilized both dSPM and eLORETA as our inverse methods to achieve cortical power estimations. These analyses resulted in high-resolution cortical power maps, with a total of 20,484 spatial points.

### Functional connectivity analysis

In this analysis, we adopted a distributed source estimation approach, focusing on the cortical surface using both dSPM and eLORETA methods and the same standard fsaverage source template we used for surface estimation. The inverse solution involved creating an inverse operator with the forward solution and noise covariance model (calculated from the selected baseline segments of each raw data), setting the loose parameter to 0.2 and the depth parameter to 0.8, to balance localization accuracy and depth weighting. We then computed the inverse solution with an signal-to-noise ratio (SNR) of 3.0, applying dSPM and eLORETA with the orientation set to normal, targeting the component perpendicular to the cortical surface. Subsequently, we partitioned the source coordinates into 68 cortical labels (34 per hemisphere) using FreeSurfer (60) aparc (61) cortical parcellation, calculating the time courses for these labels in mean-flip mode to minimize signal cancellation. For the connectivity analysis, we utilized both the Phase Lag Index (PLI) (62) and the debiased weighted Phase Lag Index (wPLI debiased) (63) to measure phase synchronization, thereby enhancing our investigation of spectral connectivity. This analysis was conducted through a multitaper approach, incorporating adaptive multitapering to improve the accuracy of spectral estimation. In the absence of a predefined adjacency sparse matrix to use for cluster permutation testing, we used a radius neighbors graph (79) algorithm to define parcel adjacencies. This algorithm constructs a graph based on the neighborhood radius around each parcel.

### Oscillatory activation extent of lucidity

In addition to our contrasts-based analyses, we focused on the LD segment to explore the spatiotemporal extent of spectral activities concomitant with LD that extend to both sides of the initial eye signaling. To examine activations at both sensor and source levels, we selected high-density EEG data (64 common channels from 19 subjects). We first identified sensor-level time intervals containing significant activations, which then informed search of these activations at the volumetric source level.

#### Sensor-level global field power around eye signaling onset

For each subject, we extracted a 30-second temporal window centered around the initial lucidity eye signal (LRLR), using the first 5 seconds as a baseline. We derived temporal power fluctuation patterns via computing absolute global field power (GFP) in this window for each channel and frequency band, normalizing the 25-second GFP patterns to this baseline with z-score standardization. To increase the sensitivity of detecting the potential significant time intervals of GFP fluctuations, we further applied downsampling to the GFP data by factor of 5. This adjustment aimed to reduce the complexity of the temporal domain, thereby enhancing the computational efficiency of the spatio-temporal permutation cluster test. Additionally, in the context of a small sample size (n=19), downsampling contributed to a more robust statistical analysis by diminishing the impact of temporal noise and reducing the risk of type I errors, ultimately making it easier to identify periods of significant neural activity. Since GFP represents absolute values, sensitivity to detect any activation pattern regardless of polarity is enhanced. Using region of interest (ROI) separation, we also analyzed GFP data from channels in frontal left, frontal right, central left, central right, parietal left, and parietal right regions separately within each frequency range (see Table S1). ROIs defined with specific channels: Frontal Left (Fp1, AF7, AF3, F7, F5, F3, F1, FT7, FC5, FC3, FC1), Frontal Right (Fp2, AF4, AF8, F2, F4, F6, F8, FC2, FC4, FC6, FT8), Central Left (T7, C5, C3, C1, TP7, CP5, CP3, CP1), Central Right (C2, C4, C6, T8, CP2, CP4, CP6, TP8), Parietal Left (P7, P5, P3, P1, PO7, PO3, O1), and Parietal Right (P2, P4, P6, P8, PO4, PO8, O2). These 25-second GFP data were normalized to the baseline as well. To determine the common extent of lucidity within the 25-second interval of each frequency band, we performed spatiotemporal cluster-based permutation tests on preprocessed EEG data.

#### Source-level decomposition of lucidity activation

We analyzed source-level activity within frequency bands that corresponded to time intervals of significant sensor-level activations. To accomplish this, we utilized dSPM and eLORETA for surface-based source reconstruction, focusing on cortical estimations for both the 5-second baseline and significant GFP activation segments. The normalization of these estimates involved subtracting baseline values from activation segments, followed by division by the maximum absolute value of the contrasted data, thus scaling the differences within a normalized range.

### Statistical Analyses

Statistical verification of SP cleaning across three different EEG setups (with 6, 64, and 128 channels) involved comparing the averaged short-window (600 ms) time-frequency power (dB) evoked by SPs at the beginning (i.e. raw data) and end of the pipeline (i.e. PREP followed by ASR followed by SSP). We selected channel F3 for the 6-channel setup and F7 for both the 64 and 128-channel setups. Our null hypothesis posited that there would be no significant increase in SP power in the raw data compared to the preprocessed data. We used the Shapiro-Wilk test to check normal distributions of each condition. If no normal distributions can be shown, we would use the Wilcoxon signed-rank test, and Student’s paired t-test otherwise.

In all further tests, we included “condition” (LD, E. REM, L. REM, waking) and “frequency band” (delta: 2-4 Hz, theta: 4-8 Hz, alpha: 8-12 Hz, beta: 12-30 Hz, gamma1: 30-36 Hz, gamma2: 36-45 Hz) as independent variables. For low-density analyses (regarding power spectral density, entropy and complexity), we utilized linear mixed models (64). This approach was particularly relevant given the presence of a few repeated measures within our subjects. To mitigate the risk of type 2 errors, we applied a parametric bootstrap method with 10,000 sample estimations, integrating subject IDs as random effect grouping factors. Subsequently, we performed post-hoc pairwise comparisons which were adjusted for multiple comparisons using the Bonferroni correction method. For high-density (topographical) analyses, we constrained statistical testing via paired 1-sample cluster permutation tests (65) with two-tailed approach. We applied threshold-free cluster enhancement (TFCE) (66) to inductively determine cluster thresholds (start/step of 2/0.2, 1.4/0.02, 2/0.2 for surface-based power estimation of a priori intervals, functional connectivity analyses, and GFP-guided surface reconstruction, respectively), except for sensor-level power analyses and spatio-temporal clustering test on GFP, for which we set the thresholds of 2.878 and 2.095 based on the T-distribution Percent Point Function (PPF) for alpha values less than 0.025. Furthermore, for sensor-level power, we used a step-down function of 0.001 and 10,000 permutations, whereas for all other analyses, 1000 permutations and no step-down were applied. While TFCE enhances our capacity to detect true effects by integrating both the spatial extent and the intensity of signals—thereby minimizing the risk of Type I errors—in analyses conducted with a two-sided approach, we further evaluated the clusters with the adjusted significance threshold. This adjustment was specifically implemented to control the false alarm rate, thereby introducing an additional layer of Type I error correction for the selected tests. LD was contrasted with E. REM, L. REM and waking. In addition, we contrasted both non-lucid REM segments against each other, leading to a total of 4 comparisons per high-density analysis and frequency band. All the statistical analyses except those for nonparametric cluster–based tests were conducted using JASP (67) version 0.17.3 and all other statistical tests were implemented in Python 3.9 with MNE package (68).

## Results

### Saccadic spike potential detection

Cleaning saccadic spike potential artifacts (SP) represents an important step in our newly developed preprocessing pipeline, requiring a valid identification of SPs (for details, please refer to Methods and Materials). The utilization of different HEOG data transforms for SP detection on example data is shown in Figure 3A and 3B. The introduction of the Hilbert transform enhanced our ability to detect saccadic spikes by amplifying SPs and reducing the impact of non-saccadic high-frequency spike artifacts. After visual inspection of all HEOG data in pre-LD and LD intervals, and after comparing 1st, 2nd order derivatives, and 2nd order polynomial values, we determined that polynomial fitting yielded the best approximation as a regularizer of Hilbert peaks. Certain artifacts resembling SPs, but exhibiting slightly lower amplitude, appeared in between and independent of saccadic activities. Slope regularization was less prone to such false-positive SPs, which is why we opted for polynomial fitting as the regularizer for the Hilbert series of the HEOG data (see Figure 3A). In general, we found that thresholding data above the 95% percentile could optimally detect false-positives as above-threshold peaks while keeping the number of misses minimal. Often, however, the actual percentile cutoff was set much higher (see Figure 3B). The formula of the proposed SP detection algorithm is shown in Eq. 1.

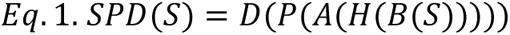

where *S* denotes the input HEOG signal, *B*(*x*) signifies the application of a bandpass filter to signal *x*, isolating the frequency components of interest, *H*(*x*) is the Hilbert transform applied to *x*, generating the envelope signal, *A*(*x*) represents taking the absolute value of *x*, *P*(*x*) is the 2nd-order polynomial fitting function, *D*(*x*) is the peak detection function, identifying the locations of SPs in the processed signal and *SPD*(*x*) is the overall saccadic potential detection algorithm. After implementing the proposed SP event detection algorithm we subjected all detected SP candidates to further visual inspection, which revealed that the algorithm had identified a few erroneous SP events, approximately 12%, which we subsequently discarded manually.

### Saccadic spike potential cleaning

Figure 3C displays an example time series of a 100-second interval around left-right-left-right eye signaling (LRLR) around one LD episode. The raw HEOG is shown in the top row, while event-related spectral perturbations (ERSPs) of progressive preprocessing steps are displayed in the rows below. From the raw data (first ERSP row), three distinct types of SP-induced artifacts are visible in the time-frequency domain: First, higher SP-correlated power can be shown above our analysis cutoff of 50 Hz, compared to the power below 50 Hz. Second, even higher SP artifact-related power in low frequency bands (below 5 Hz) can also be seen. Lastly, artifactual broadband effects are visible as vertical stripes of elevated activity. After the first preprocessing step (PREP (41); second ERSP row in Figure 3C), we see the eradication of artifactual broad-band activity and some diminishing of artifactual low-frequency activity. Since PREP is not designed to clean artifacts in EEG, this initial cleaning should be improvable by further preprocessing steps. Indeed, the addition of artifact subspace reconstruction (ASR (69); third ERSP row) visibly reduced artifactual low-frequency activity even further, but leaves some bursts of activity that are correlated with SP events (see vertical dashed lines in Figure 3C), especially between 30 and 45 Hz. Considering that up to this point only EEG-based cleaning had been performed, subsequent signal space projection (SSP) based on HEOG patterns could remove all remaining types of SP artifacts as expected.

Figure 3D shows the effects of the cumulative preprocessing steps on grand average SP-induced ERSPs of frontal channels. In both medium-(64 channels; active electrodes; 16 SP events) and high-density (128 channels; passive electrodes; 63 SP events) EEG setups, each additional preprocessing step progressively cleans phasic activities correlated with SPs but retains unrelated tonic activity, which is most apparent in the retained contrast between frequencies below and above 10 Hz. Statistical analyses support these observations: Since the values are normally distributed (Shapiro-Wilk, p = .106, .996, and .208 for 6-channel (14 SP events), 64-channel and 128-channel data, respectively), we employed t tests. The SP evoked power values are significantly lower after the final preprocessing stage for 6-channel (t(13) = 3.012; p = .005; d = 0.805), 64-channel (t(15) = 10.319; p < .001; d = 2.580) and 128-channel setups (t(62) = 2.971; p = .002; d = 0.374).

Visually, the application of ASR on top of PREP exhibits the most drastic changes in artifact removal (compare second and third rows in Figure 3D). Note, however, that right-hemispheric artifacts in high-density passive setups appear more resistant to cleaning (rightmost column in Figure 3D). This pattern is also reflected in the ∼40 Hz-band topographies in Figure 3E (bottom row): Increased cleaning along the preprocessing pipeline is shown with the addition of ASR visibly decreasing the most artifacts. Again, some right-hemispheric artifactual components are retained even after SSP (see Figure 3E, right side). This might be due to different datasets having systematically different placement of HEOG electrodes in a way that rightward saccades could not be detected as easily by our algorithm and SSP. GFP (gray areas in Figure 3E, upper row) again confirm that the combination of PREP and ASR have the biggest effect on SP-related activity reduction. The same is shown for Hilbert envelopes across EEG channels (colored lines in Figure 3E, upper row). Note that in 64-channel active setup data, fluctuations visibly oscillate more often (indicative of previous and upcoming saccades), more slowly and with higher amplitude than in 128-channel passive setups. This might be because instructions for participants from different datasets were systematically different. For example, in 64-channel datasets participants might have been instructed to do extensive LRLR signals, which take longer to perform, and to perform these signals multiple times in a row. Crucially, note that all these oscillations of saccadic origin are resolved after applying PREP and ASR, which highlights ASR in handling non-stationary artifacts.

### Power spectral density analysis results

With the validity of our preprocessing demonstrated, the following analyses will focus on exploring electrophysiological correlates of LD. In all analyses, we investigated the following within-subject and within-session conditions: waking, LD, earlier non-lucid REM sleep (E. REM), and later non-lucid REM sleep (L. REM, i.e. similar times as LD periods; see Figure 1A; see Materials and Methods for further details). Results of global, sensor-level power analyses are shown in Figure 4A-C. As expected, the waking condition showed a pronounced power increase in the alpha band (8-12 Hz) compared to non-waking stages. A robust increase in power can also be seen from 20 Hz upwards (Figure 4A) during waking.

**Figure 4.**
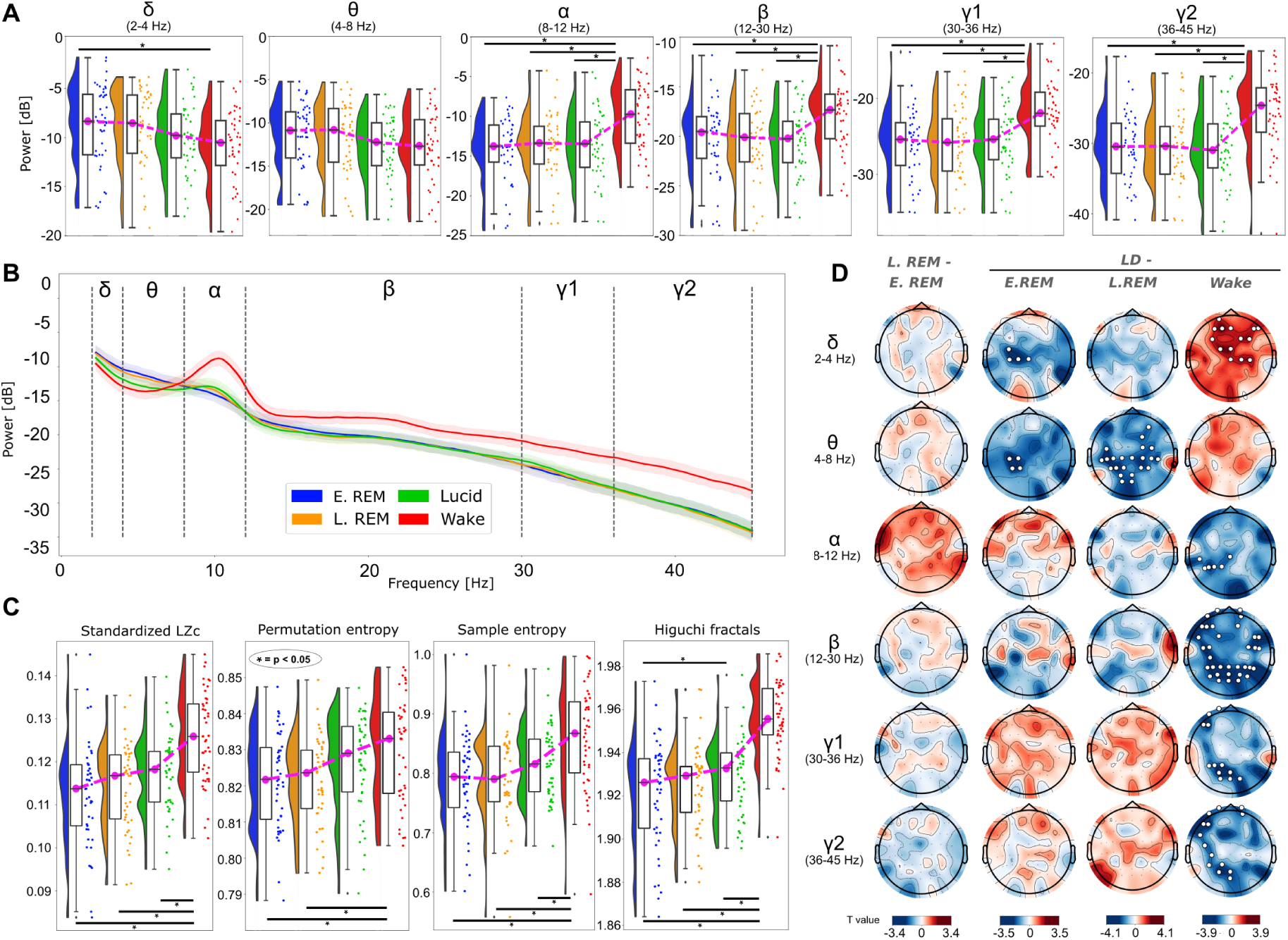
Sensor-level PSD, complexity, entropy and topographical power analysis results. **A:** Statistical comparison of PSDs across conditions for each frequency band. The colored shapes represent kernel density estimation (KDEs) of the data distribution. Overlaid box plots display the median and interquartile range, with whiskers indicating the data spread. The magenta line connects the mean values across conditions. **B:** Grand average PSDs by across 6-common channels with shaded areas showing 95% confidence intervals. Error bars show standard error of the mean. **C:** Statistical comparison of LZc, entropy measures, and HF across conditions. KDEs are shown in color to represent the data distribution, complemented by box plots that indicate the median and interquartile range. **D:** Sensor-level topography maps contrasting tonic power in LD, REM and waking conditions within each frequency band. Colors show T values representing relative changes in power. White dots point to electrode locations that belong to significant channel clusters. PSD: power spectral density; LZc: Lempel-Ziv complexity; HF: Higuchi fractals; LD: lucid dreaming; REM: rapid eye movement sleep

Frequency-band-specific linear mixed models (LMM) analyses revealed significant differences between conditions in the delta (p = .029), alpha (p = .001), beta (p = .010), gamma1 (p = .007), and gamma2 frequency bands (p = .001). The theta band failed to reach statistical significance (p = .259). Post-hoc analyses revealed that the waking condition drives the observed effects because waking power differs from all other conditions (see Figure 4C): In the delta band only waking power (-10.657 ± 0.738) was higher than E. REM (-8.372 ± 0.799); in the alpha band, waking power (-9.493 ± 0.746) was higher than E. REM (-13.529 ± 0.773) (p < .001), L. REM (- 13.036 ± 0.824) (p < .001) and LD (-12.749 ± 0.859) (p < .001); in the beta band, waking power (-17.799 ± 0.706) was higher than E. REM (-20.154 ± 0.748) (p = .017), L. REM (-20.352 ± 0.748) (p = .011) and LD (-20.348 ± 0.743) (p = .011); in the gamma1 band, waking power (-21.555 ± 0.768) was higher than E. REM (-25.228 ± 0.935) (p < .001), L. REM (-25.063 ± 0.955) (p = .001) and LD (-24.830 ± 0.913) (p = .002); and in the gamma2 band, waking power (-24.832 ± 0.769) was higher than E. REM (-29.341 ± 1.014) (p < .001), L. REM (-29.434 ± 1.058) (p < .001) and LD (-29.338 ± 1.022) (p < .001). Notably, no statistical differences were observed within REM variations (E. REM and L. REM) and between the REM variations and LD in any frequency band.

### Entropy, Lempel-ziv complexity and Fractal analyzes results

Markers of Lempel-ziv complexity (LZc) (70), sample entropy (SE) (55), permutation entropy (PE) (71) and Higuchi fractal dimension (HF) (57) have been shown to covary with conscious awareness (53, 72, 73), which makes them potential candidates for metrically capturing lucidity (18). The differences between waking, LD and non-lucid REM sleep with regards to entropy and complexity markers are shown in Figure 4C. LMM results reveal that alll measures differ significantly between conditions (LZc p < .001; PE p = .004; SE p < .001; HF p < .001) . Post hoc analyses reveal that the waking condition drives these effects. Specifically, LZc values are significantly higher in the waking condition (0.126 ± 0.002) compared to early REM (0.113 ± 0.002) (p < .001), late REM (0.114 ± 0.002) (p < .001), and LD (0.117 ± 0.002) (p < .001). We found a similar pattern in SE values: waking values (0.859 ± 0.011) are significantly higher than those of early REM (0.790 ± 0.011) (p < .001), late REM (0.794 ± 0.011) (p < .001), and LD (0.817 ± 0.011) (p = .002). Regarding PE, waking levels (0.828 ± 0.002) were significantly higher only compared to early REM (0.821 ± 0.002) (p = .024) and late REM (0.820 ± 0.002) (p = .012). Interestingly, we found no significant difference between waking and LD. As our final marker, the HF showed the most significant pattern (LMM p < .001) when compared to other markers: waking values (1.957 ± 0.003) are significantly higher than those of early REM (1.919 ± 0.005) (p < .001), late REM (1.923 ± 0.004) (p < .001), and LD (1.929 ± 0.004) (p < .001). Additionally, we found that LD is significantly higher than early REM (p = .044). For all markers, no statistically significant difference could be shown between LD and non-lucid REM conditions.

### Topographical power analysis results

A sufficient sample size of high-density EEG recordings allowed us to explore channel clusters showing significant differences between conditions across frequency bands of interest (see Figure 4D). Firstly, we observed no clusters that could significantly distinguish between L. REM and E. REM in any of the frequency bands. However, when contrasting LD with waking, we note significant topographical differences in all frequency bands but theta. Specifically, in the delta band, we observed higher power in the frontal and central regions during LD; in the alpha band, a significant decrease in the left-occipital region; in the beta band, a near-global decrease except the central region; in the gamma1 band, significant decreases in the frontal-left and occipital regions; and in the gamma2 band a significant power drop particularly in the frontal-left and extending to the frontal and left-parietal and occipital regions. Sensor-level topography of theta band power differs significantly between LD and non-lucid REM (i.e., both E. REM and L. REM) with an observed theta decrease in LD compared with L. REM over central and extending to parietal and occipital channels, and an observed theta decrease in LD compared with E. REM in the central-left region. We also found significant differences in LD delta band power compared to E. REM delta band power, with stronger power differences over the central-left channels, but no differences in LD delta compared to L. REM delta.

### Surface-based source reconstruction results

To further investigate neural activation distinguishing LD from waking and non-lucid dreaming we employed source reconstruction. To benefit from its excellent noise suppression characteristics, we planned to use beamforming approaches (74, 75), however, given that we are combining EEG data with template MRIs, there are concerns whether our forward models would be of sufficient quality for beamforming (76). Thus, we compared the beamforming output to two more robust source reconstruction approaches, namely (a) dynamic statistical parametric mapping (dSPM) (58) and (b) exact low-resolution electromagnetic tomography (eLORETA) (59). This comparison consolidated our concerns and thus we only report the results of the two minimum norm estimation approaches but not the beamforming. Grand average distributions of cortical power per conditions and frequency band are depicted in Figure 5. Results of statistical comparisons are shown in Figure 6A for dSPM and eLORETA, respectively. Comparing the two methodologies, we note substantial overlap in the findings of the two methods as well as with sensor-level topographical results (see Figure 4D) in frequency bands from 8 Hz to 45 Hz, underscoring the robustness of our findings. Specifically, within the alpha, beta, and gamma frequency ranges, we find power reductions in LD relative to waking conditions. Both methodologies identified alpha power reductions primarily in the occipital region, whereas beta power showed a more extensive decrease spanning the bilateral temporal, frontal, and occipital lobes. In the gamma frequency bands, eLORETA indicated a widespread cortical power decline with notable decreases in the left and particularly the right fronto-lateral regions, while dSPM suggested a more localized reduction, especially in the right frontal region for gamma2. Furthermore, the findings revealed localized significant variations between LD and L. REM sleep for both dSPM and eLORETA. dSPM pointed to a theta decrease in the right occipital region with extension from posterior aspects towards the parietal lobe encompassing precuneus in LD, a pattern similar to the beta reduction observed mainly in the precuneus with traces in the occipital area. eLORETA identified beta power reductions as well, but showing a more distributed profile. Additionally, an increase in gamma1 power during LD in the left fronto-lateral region as compared to L.REM was observed via dSPM only. No significant differences in cortical surface power were observed between LD and E. REM, or between L. REM and E. REM, across all frequency bands. Significant connections employing weighted debiased PLI measures are presented Figure 7.

**Figure 5.**
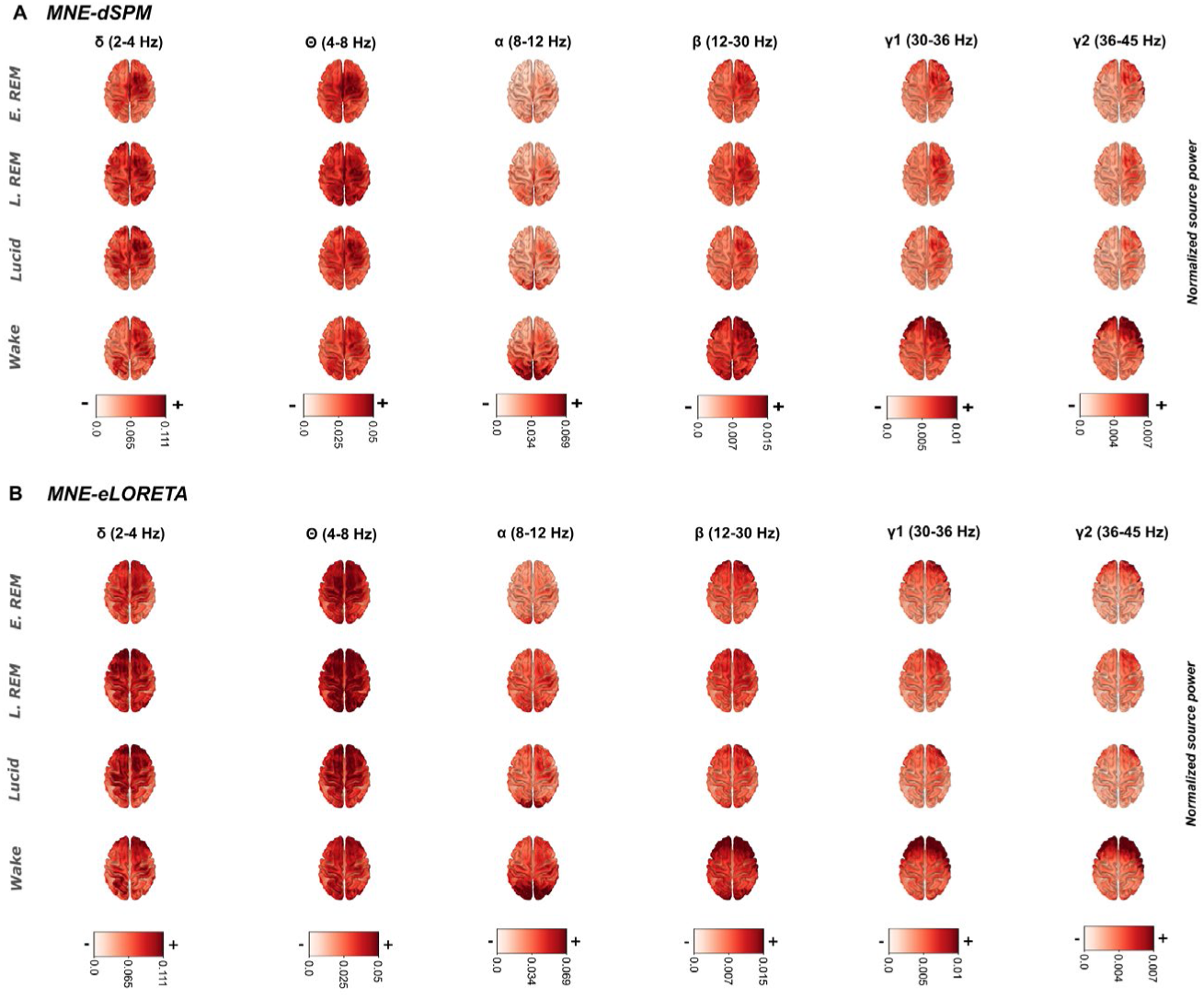
Normalized source-level power across frequency bands and conditions, utilizing two methodologies: MNE-dSPM (Figure part A) and MNE-eLORETA (Figure part B). MNE: minimum norm estimation; dSPM: dynamic statistical parametric mapping; eLORETA:exact low-resolution electromagnetic tomography.

**Figure 6.**
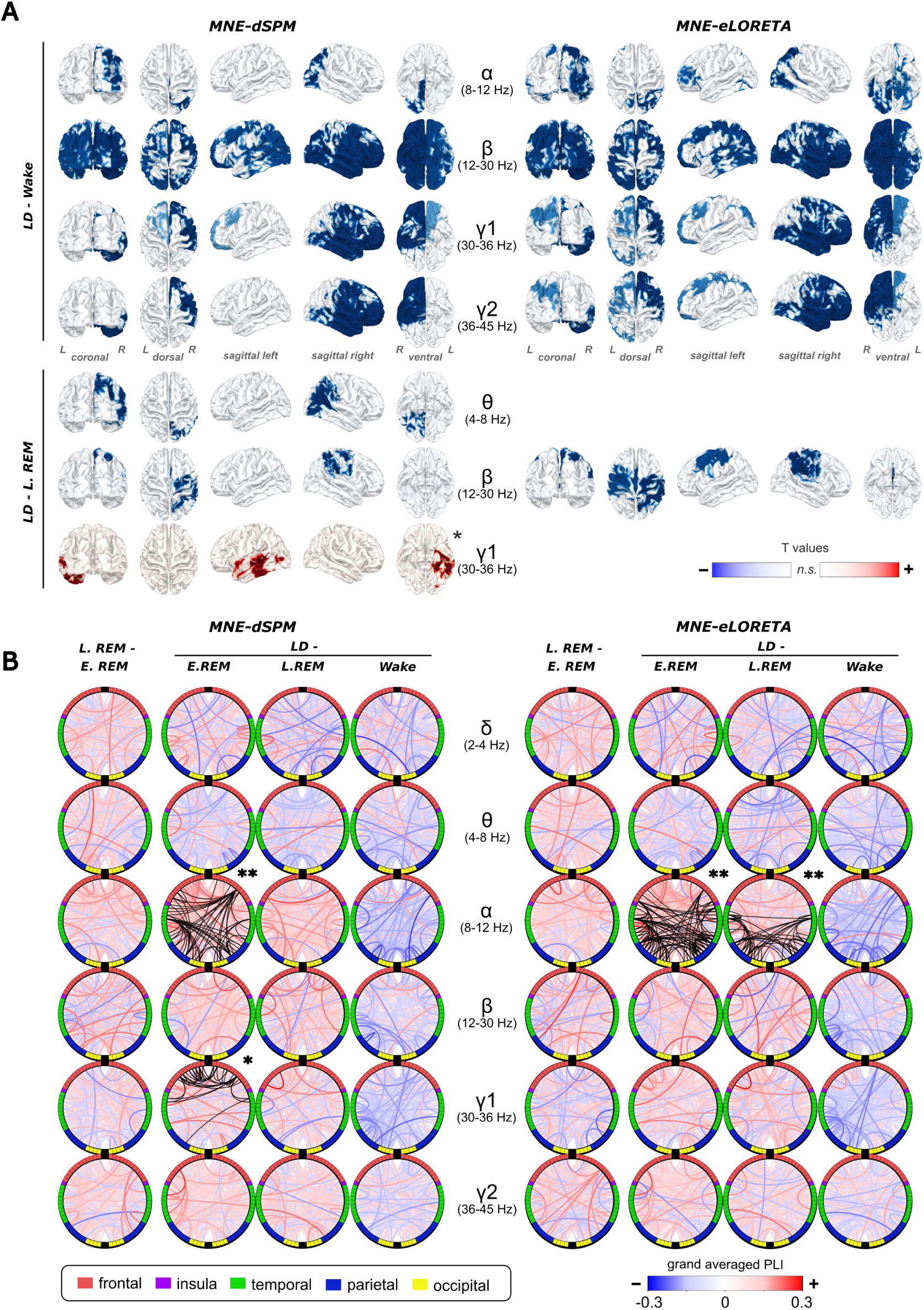
**A:** Thresholded surface-based source-level power for each frequency band derived from dSPM and eLORETA and plotted over cortex images. Each surface significance is shown with 5 different views (coronal, dorsal, left sagittal, right sagittal, and ventral). Power values of statistically significant clusters contrasting LD, REM and waking for both dSPM and eLORETA methods. All shown clusters were significant with p < .025, except clusters marked with *, which were significant with p < .05. The label “n.s” denotes regions where the results were not statistically significant. B: Grand average contrasted connectivities between L. REM and E.REM, LD and E. REM, LD and L. REM, and LD and wake across frequency ranges, analyzed using dSPM and eLORETA. Increases in PLI connectivity patterns, significant through cluster-permutation tests, are highlighted in black color (dSPM: LD - E. REM alpha and gamma1; eLORETA: LD - E. REM and LD - L. REM alpha), with no significant reductions detected. All statistical tests were conducted with and without additional bilateral significance threshold adjustments, with the symbols “*” and “**” indicating the p-threshold (.025 and .05, respectively) for each visualized significance test. LD: lucid dreaming; REM: rapid eye movement sleep; PLI: phase lag index

**Figure 7.**
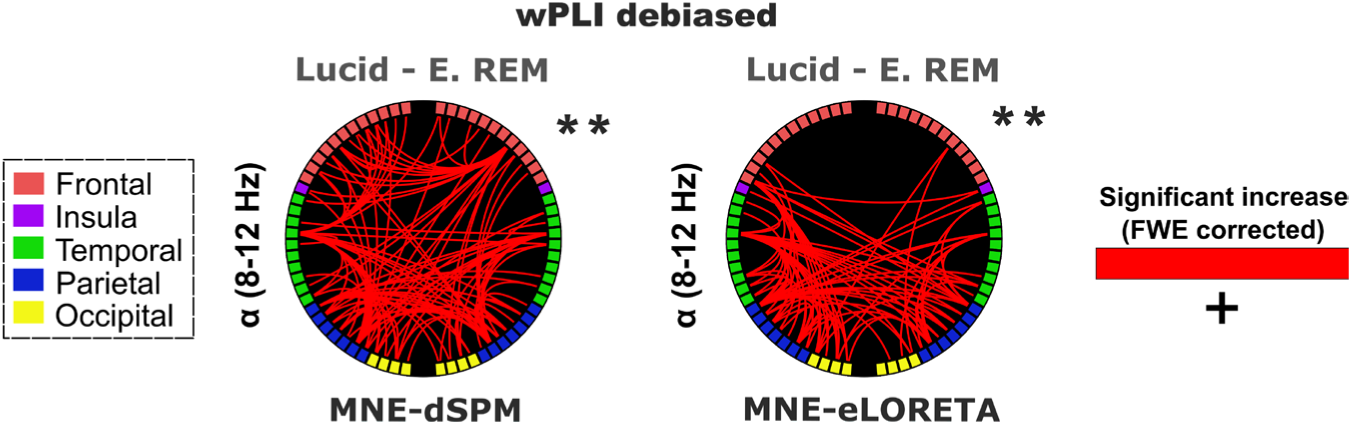
Normalized source-level connectivity across frequency bands and conditions, utilizing two methodologies: MNE-dSPM (upper part) and MNE-eLORETA (lower part). Left side: grand average PLI values per frequency band and condition. Right side: significant connections using debiased wPLI. Two asterisks mark significance with p < .025. MNE: minimum norm estimation; dSPM: dynamic statistical parametric mapping; eLORETA:exact low-resolution electromagnetic tomography; PLI: phase lag index; wPLI: weighted phase lag index.

### Functional connectivity results

Cortical measures of functional connectivity allowed us to explore correlates of LD even deeper. Grand average phase lag indices (PLIs; see Figure 6B) reveal robust increases in alpha connectivity during LD as compared with E. REM across analysis methods (i.e., across dSPM, eLORETA with PLI/wPLI debiased approaches). Inspecting shared connections across all methods, robust hub areas become evident, most prominently the left superior temporal gyrus, but also right superior parietal gyrus, right superior frontal gyrus and left lingual gyrus (see Figure 6B). Significantly increased alpha connectivity during LD could be shown against L. REM as well, however only in PLI measures and when using eLORETA (see Figure 6B). Frontal regions were not identified in this connectivity pattern, which overlaps with the auditory resting state network and partly the visual and posterior parts of the default mode network as well. Apart from alpha, we noted increased gamma-1 band connectivity during LD as compared to E. REM when employing PLI measures from dSPM, almost exclusively in frontotemporal regions. No other significant connectivity effects could be shown. Specifically, we observed no differences between LD and waking or between temporally distant non-lucid REM intervals.

### Temporal dynamics of lucidity activation

In a last analysis, we tested the temporal dynamics of LD around the initial LRLR eye signaling (within a symmetrical search window of 30 seconds) via spatiotemporal clustering of global field potentials (GFPs). The first 5 seconds of the search window served as the baseline (for further details, refer to the Materials and Methods section). All activation patterns are shown in Figure 8. According to the results of the spatio-temporal cluster tests, we found significant activation in the alpha band, which exceeded the cluster threshold in between 1.75 seconds before and 3.25 seconds after LRLR onset. However, no significant differences were identified at the source level when contrasted with the baseline in this band. We also detected significant activation patterns at both the sensor and source levels in the gamma1 and gamma2 bands. Specifically, in the gamma1 band, spatiotemporal clusters can be observed between 2 seconds before and 7 seconds after initial LRLR, leading to sensor-level activation primarily in the occipital-left region, as determined by the topographic layout of GFP activations. In the gamma1 frequency range, both MNE-dSPM and eLORETA methods identified marked increases in cortical activity. MNE-dSPM reveals bilateral frontal lobe engagement, with notable extensions into the central and parietal regions, alongside a significant activation in the left temporal lobe. eLORETA echoes these findings but with more pronounced and extensive activations across the frontal and parietal lobes, notably highlighting the right precuneus, and enveloping the temporal lobes bilaterally. Additionally, in the gamma2 frequency band, significant sensor-level activation was detected that exceeded cluster thresholds between 1 second before and 5 seconds after initial LRLR, particularly over occipital-left channels. When examining the same time interval at the source level, robust activations were discerned at the sensor level that surpassed cluster significance thresholds within the period extending from one second prior to five seconds following the onset of LRLR eye signals, predominantly over the occipital-left channels. This sensor-level activity is mirrored at the source level with predominant activations in the left fronto-lateral and left temporal regions as delineated by both dSPM and eLORETA analyses.

**Figure 8.**
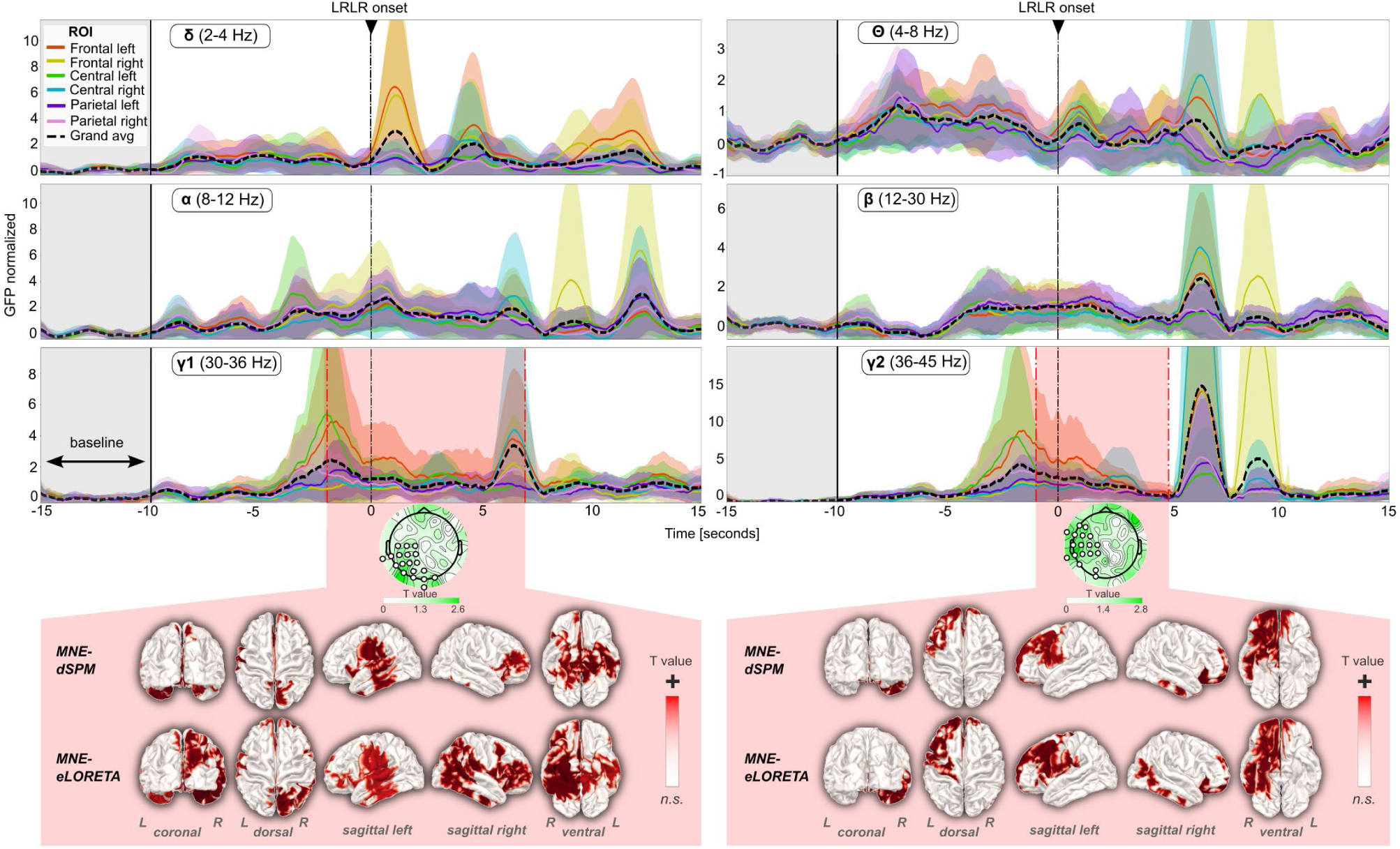
ROI-and frequency-band specific GFP fluctuations of EEG data during LD around the time point of LRLR onset. Error shadings represent 95% confidence intervals. GFPs are normalized on the baseline interval which is shown in light red shading 15 to 10 seconds before LRLR onset. Time intervals of significant spatio-temporal activation of EEG data are marked in gray shading and with a dash-dot line. Inlays show clusters exceeding the cluster threshold at the sensor-and source-level of this time area. The label “n.s” denotes regions where the results were not statistically significant. ROI: region of interest; GFP: global field power; LRLR: left-right-left-right eye signal.

## Discussion

Lucid dreaming (LD) is an intriguing state of consciousness that remained poorly understood at the electrophysiological level, largely due to low statistical power, inadequate saccadic artifact correction, and limited sensitivity to source-level activities. Here we employed thorough saccadic potential cleaning and cortical source reconstruction in the highest sample size with most densely sampled topographical EEG resolution in the LD field to date. In comparison to non-lucid REM sleep, we observed a reduction in posterior theta power, and a robust beta power decrease in the right temporoparietal junction. Besides, we found gamma activity involving the precuneus, left prefrontal cortex, and frontopolar areas around the time of the first LD eye signaling. Finally, we identified enhanced connectivity in the alpha band with a prominent hub in the left superior temporal gyrus, as well as increased gamma connectivity within bilateral frontotemporal regions, as discussed below.

Since our preprocessing pipeline was inclusive of low-density datasets, we could revisit earlier sensor-level spectral findings (for an overview, see (17)) with a well-powered sample size. We found that LD significantly differed from the waking state across most frequency bands (Figure 4A) with LD showing systematically higher power in low frequencies (2-4 Hz) and lower power in higher frequencies (8-45 Hz), a finding that aligns with established patterns of REM sleep in general (18, 51). This result also confirms that subjects are not awake, a common critique in the first years of LD research. No differences were found in the theta band (4-8 Hz), but the full spectrum (Figure 4B) shows that this may be an averaging effect over frequencies, as pronounced differences are visible approximately between 3 and 5.5 Hz. In this range, spectral power of LD is located in between the waking state and non-lucid REM sleep. Visual inspection thus suggests that traditional frequency band limits might not be sensitive enough to capture the true characteristics of LD, echoing recent calls for more inductive approaches to analyze spectral data (78). Furthermore, it has been recently shown that measures of EEG signal complexity and entropy could distinguish lucid from non-lucid REM sleep (18). With a notably larger sample size in the current study, we failed to replicate this finding with regards to signal complexity, entropy or fractals (see Figure 4C). A potential reason for this might be too much inherent heterogeneity in our data. Alternatively, a false positive result might have been obtained by chance in (18), given the multitude of potential aperiodic signal measures. A systematic investigation of these measures in larger samples of LD data would be a worthwhile future endeavor.

Investigating high-density topographies, we found frontocentral and parietal clusters with theta power reduction in LD epochs compared to REM sleep epochs (Figure 4D). This might reflect attentional processes, considering that neurofeedback protocols aimed at increasing attentional control in clinical samples are based on decreasing theta power (79, 80). Therefore, lower theta power might help in becoming lucid during REM sleep. Note that our dSPM analysis also supports this finding (Figure 6A) which localized effects mainly to right occipital areas. While (18) found widespread reductions in delta activity during lucid compared to non-lucid REM sleep, our study could only partially corroborate this result: Such a sensor-level reduction was only found when compared to earlier REM sleep. This might be an interactive effect of sleep duration and dream lucidity given that cluster tests did not find this power drop compared to later REM sleep, when the temporal proximity is much higher compared to the early REM state (Figures 1A and 4D). Note that resolving delta activity at the source level could not identify the relative power increase in LD compared to the waking state, which was shown with sensor-level analysis (Figure 6A). This might indicate that the spatial distribution of delta activity during LD shares similarities with the waking state. We further noted a significant source-level increase in gamma1 power (30-36 Hz) when contrasting lucid against later REM sleep (Figure 4D), which however was only shown via dSPM-based source reconstruction and could not be corroborated with sensor-level data (Figure 6A). This likely stems from dSPM’s noise normalization technique, which boosts signal-to-noise ratios to reveal localized neural activity not detectable with eLORETA and sensor-level analyses (58). Gamma1 activity increases were localized primarily to left-hemispheric temporal areas and may accompany processes of verbal insight (81). Furthermore, increased activity within the middle temporal gyrus during LD has been hypothesized within a framework of predictive coding and resolution of low-level sensory prediction errors (82).

A consistent finding from both cortical source localization techniques (MNE-dSPM and MNE-eLORETA) is the reduction of beta band activity in LD relative to temporally close non-lucid REM sleep (Figure 6A). This effect was not observed at the sensor level, suggesting beta reduction during LD might only be discernible with the refined spatial resolution MNE techniques offer. Beta band activity is theorized to underlie the maintenance of a cognitive status quo (83). Its reduction therefore aligns well with a key requirement for LD, which is the reassessment of the veridicality of currently perceived reality. However, this assumption awaits confirmation from future research. Note that dSPM’s noise normalization approach provides more focal source clusters, revealing the temporo-parietal junction (TPJ) in the right hemisphere as a likely origin of beta activity. According to a recent theoretical framework (82), LD puts higher precision weighting on lower sensory predictions, and increased prediction error resolution would result in heightened TPJ activity during LD. It has been found that the TPJ integrates visual, auditory, tactile, proprioceptive, and vestibular information, contributing to self-consciousness and internal body imagery (84). Disrupting TPJ activity during the waking state with magnetic (85) or electric (86) transcranial stimulation can cause an out-of-body experience, defined as a subjective sensation of being “outside the own body” (87). Interestingly, LD and out-of-body experiences have been linked (88–90). In our analysis of neurophysiological correlates around the initial lucidity eye signals, we observed no such beta effects (Figure 8), potentially due to the brief baseline period not adequately capturing the sustained nature of beta-mediated metacognition (83).

Preprocessing data with our pipeline supports recent findings (18) that LD is not characterized by EEG frontal activity in frequencies around 40 Hz when SP artifacts are sensibly cleaned (Figures 3E and 3D). However, theoretically this does not exclude the possibility that frontolateral gamma plays a role in LD (21, 91), but is overshadowed by saccadic potentials. Gamma oscillations are associated with attention, sensory integration, and consciousness (92–94) — processes that are engaged during LD tasks such as flying and visual exploration. We observed gamma activations around the initial eye signal marking lucidity’s onset, suggesting it might precede epochs marked as lucid via eye signaling (Figure 8). Source-level analysis revealed an increase in gamma1 (30-36 Hz) activity around the initial eye signaling, which might represent active metacognitive processing related to the shift from non-lucid dreaming to conscious awareness. Importantly, source localization resolved this gamma1 activity partly to the precuneus in the right hemisphere. The precuneus facilitates complex cognitive functions through gamma oscillations, particularly during states of heightened conscious awareness (95). Precuneus activity is also significantly modulated during self-related cognition (96) as well as closed-eyes waking imagery induced by psychedelics such as Ayahuasca (97) and LSD (98), which have remarkable similarities with dreaming in general (99–101) and LD in particular (102). This makes the precuneus a likely neuroanatomical correlate of LD, which is in line with earlier fMRI findings showing the precuneus as one of the most prominently activated areas in LD compared with non-lucid REM dreaming (103). We further observed increases in gamma2 (36-45 Hz) related to the initial eye signaling, most prominently in left prefrontal and frontopolar cortices. This localization, together with the fact that gamma2 includes 40 Hz that is notoriously difficult to clean from eye-movement-related artifacts (18, 29, 30), alerts to a likely artifactual origin of this finding. Disentangling the exact contributions of such non-neural components from true neural sources requires imaging methods that are not sensitive to electric currents, such as fMRI. While previous MRI research identified frontopolar cortices to be associated with LD (103–105), and the preprocessing pipeline tested in this study satisfactorily removed saccadic traces in EEG (Figure 3D and 3E), it remains doubtful if localized neural gamma oscillations are involved in the present finding. Given the high-amplitude EOG from conscious eye signaling that is necessarily included in the analysis interval, it is more likely that our study supports the recent finding by Baird and colleagues (18) that saccadic activity explains frontal 40-Hz signals found to be associated with LD (21), via source reconstruction methods as well.

The spectral connectivity results point to increased long-range alpha communication during LD as compared to non-lucid REM sleep, especially within the left-hemispheric auditory resting state network (Figure 6B). Statistically, this effect is mostly detected in systematically earlier non-lucid REM episodes, alerting to a potential confound of sleep duration. Across source localization and connectivity computation methods, we note alpha connectivity between right superior frontal gyrus and left lingual gyrus. The involvement of anterior frontal cortices in LD is noteworthy given that during non-lucid REM sleep, the lateral prefrontal and frontopolar cortices are downregulated (106) and uncoupled from posterior regions, effectively separating executive functions from sensory perception (107, 108). This overall pattern could underlie the lack of judgment and passive acceptance of bizarreness during ordinary, non-lucid dreaming (109). Alpha connectivity increase stands in contrast with psychedelics-induced decreases in alpha connectivity (110), even though psychedelic experiences share with LD experiences many subjective features such as vivid visual imagery, enhanced emotional intensity and increased metacognition (111). While psychedelics often lead to a dissolution of ego and decreased self-referential processing within the default mode network (DMN), LD may actually harness elements of self-awareness and control that require enhanced alpha connectivity. Similar to psychedelics, the alpha reduction seen during trance states (112) induced by shamanic rituals suggest an altered state of consciousness divergent from the self-directed focus experienced in LD.

Regarding other limitations of our study, it should be noted that subjective aspects beyond dream awareness could also influence the differences observed in LD, such as the often-reported increase in perceptual vividness, especially visual (33). Unfortunately, the lack of non-lucid dream reports precludes a statistical control for sleep mentation, and future studies should include such reports. Additionally, controlling for dream content might yield more robust results, as the representation of background lucidity in the signal may be too weak when mixed with varied dream content. The moment of lucid insight might serve as a unifying event that is highly similar across different lucid dreams and individuals, potentially offering meaningful insights into the mechanisms of LD. While capturing this event is methodologically very challenging, we consider it a promising avenue for future research.

To this end, the tonic brain activity identified in our study might serve as a guide to better define the temporal extent of individual lucid dreams. Furthermore, despite the advancements our data cleaning pipeline allows, the potential for overcleaning and inadvertently diminishing signals of interest is a risk that can never be entirely eliminated. Still, while artifacts were cleaned by our preprocessing protocol (Figure 3), signals of interest are also retained as reflected in the spectral characteristics (Figure 4A and B), which highlight the pronounced differences between the waking state and REM sleep (including LD). These differences comprise a wake alpha peak near-absent in sleep (113) and relatively increased higher power in higher frequency bands during the waking state (114). Mirroring the spectral results, waking epochs also differentiated from all REM sleep epochs with regards to entropy, complexity, and fractal dimension measures, as expected (73, 93, 115, 116) (Figure 4C).

In summary, electrophysiological source localization measured from lucid REM sleep offers important insights into the nature of the brain’s ability to determine whether currently active world models are aligned with one’s environment (as in functional waking perception or LD) or not (as in non-lucid dreams and hallucinations). LD thus offers a unique avenue for voluntarily interacting with and immersively adjusting internal world models whose dysfunction is often at the root of various mental disorders (104, 105). The ability to adjust these dysfunctional models during dreaming presents a potentially useful tool for therapy (111, 117–119). The electrophysiological characterization of LD across a representative dataset shown in this work deepens our understanding of the cortical activities associated with metacognitive insight into an ongoing state of non-veridical world representation. It may also facilitate the development of neurofeedback and brain-computer interface technologies aimed at inducing LD towards unlocking its full clinical potential.

## Author Contributions

Conceptualization: ÇD, KA, MZ, SM-R, SR, MZ, NA, MD; Data collection: ÇD, JG, KA, KL, CF, CR, SM-R, DE, SL; Methodology: ÇD, BW; Data curation: ÇD, JG, KA, KL, XW, ZZ, NA; Preprocessing: ÇD; Formal analysis: ÇD; Statistical testing & visualization: ÇD; Writing -original draft: ÇD, NA; Writing -review and editing: ÇD, JG, KA, KL, CF, CR, BW, XW, ZZ, AS, DE, SL, SM-R, SR, MZ, NA, MD; Supervision: KA, MZ, AS, SL, SR, NA, MD

## Acknowledgments

We are grateful for all students and assistants involved in spending overnights and earliest mornings to help provide the database for this study and make this project a reality. We also thank Mahdad Jafarzadeh Esfahani and Pedro Reis Oliveira for assisting in sleep scoring.

